# The coordination between xylem and bark hydraulics in temperate Rosaceae species

**DOI:** 10.64898/2026.06.06.730605

**Authors:** Radek Jupa, Terezie Pátková, Jan Binter, Jiří Doležal, Michael Peter Nobis, Stefan Mayr, Vít Gloser

## Abstract

Xylem and bark properties influence tree growth and drought resistance, yet their functional coordination and their environmental drivers remain unclear.

We assessed xylem–bark coordination in branches of eight temperate woody Rosaceae species spanning different ecological preferences. We quantified xylem hydraulic efficiency and safety alongside bark traits governing permeability, hygroscopic water exchange, water storage, and anatomy, and evaluated phylogenetic signal and climatic associations.

Bark water vapor conductance (G_bark_) increased with maximum xylem hydraulic conductivity (Kh) and with xylem water potential at 50% loss of conductivity (P_50_), indicating species with more efficient but more embolism-vulnerable xylem developed more permeable bark. Species with higher G_bark_ showed reduced hygroscopic absorption time, consistent with faster rehydration from atmospheric water vapor. Both G_bark_ and P_50_ were phylogenetically conserved and covaried with climatic factors, namely air temperature, vapor pressure deficit (VPD), and isothermality. Species from warmer, high-VPD climates with greater diurnal temperature variability combined higher bark permeability with more vulnerable xylem, implying a shift from embolism avoidance to embolism tolerance strategies.

Overall, xylem and bark hydraulics in Rosaceae evolved in concert along diurnal and annual gradients of evaporative demand, showing that drought resistance in woody angiosperms cannot be understood without considering bark traits alongside xylem function.

**Graphical abstract:** 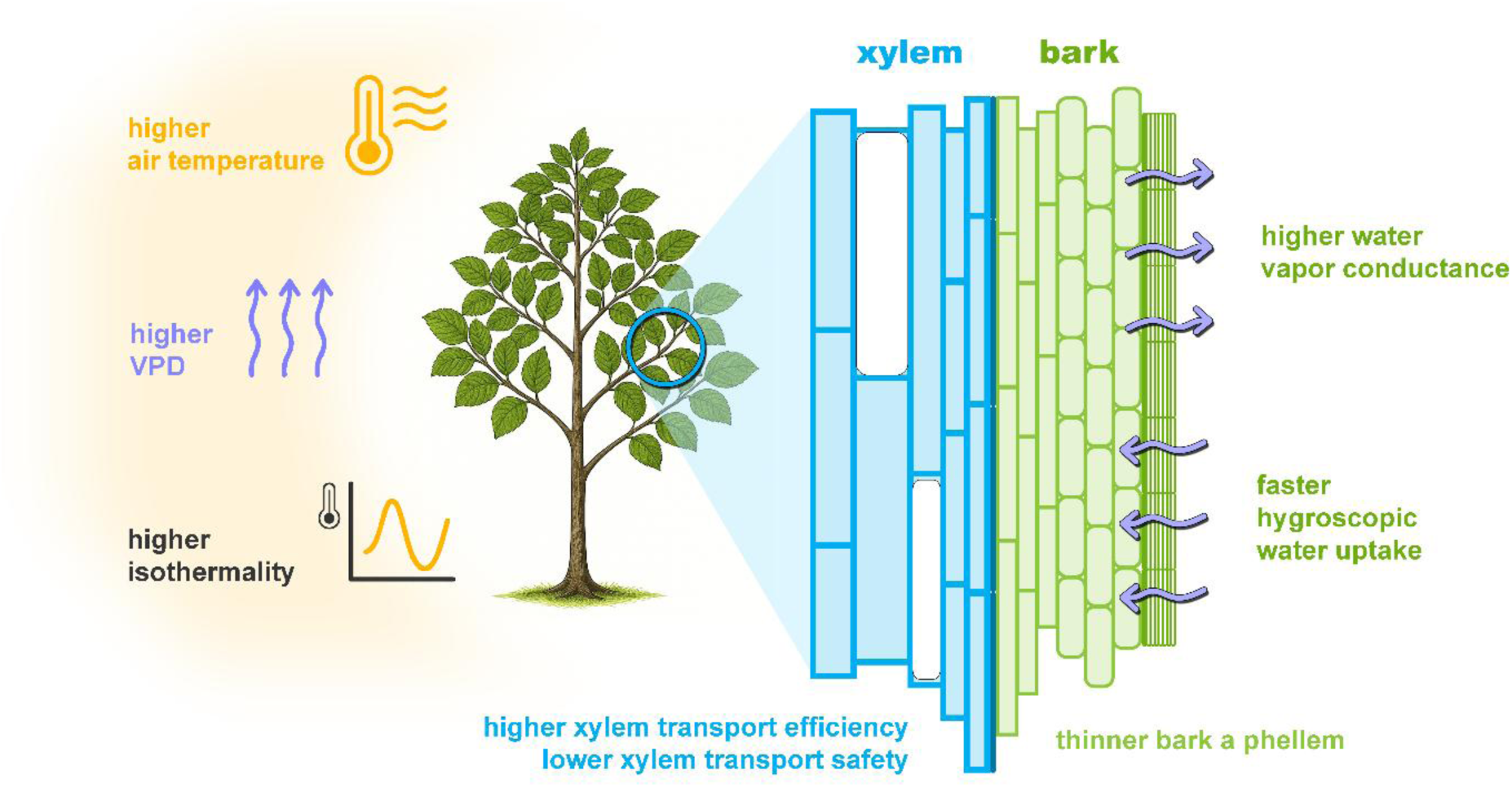

**Caption:** This study shows that xylem and bark in Rosaceae species form an integrated functional system in which xylem hydraulic safety, efficiency, and bark permeability are jointly tuned along diurnal and annual gradients of air temperature and evaporative demand.

## Introduction

Increasing drought frequency and intensity under climate change have elevated concerns about drought-induced tree mortality, driving efforts to identify the mechanistic traits that underpin drought resistance and predict species vulnerability under future climates (Adams et al., 2017; Brodribb et al., 2020; Groover et al., 2025; Chen et al., 2025). Tree drought resistance emerges from the coordinated integration of multiple structural, physiological, biochemical, and molecular mechanisms shaped over evolutionary time, which collectively regulate water uptake, transport, storage, and loss as well as the maintenance of living tissues (McDowell et al., 2008; Choat et al., 2018; McDowell et al., 2022). Among these mechanisms, xylem water transport is a central determinant of whole-tree drought resistance, governing the plant’s hydraulic integrity under drought stress (Choat et al., 2018; Mantova et al., 2022).

Xylem enables long-distance transport and distribution of water and mineral nutrients throughout the plant body, thereby underpinning tree growth and carbon assimilation (Tyree & Zimmermann, 2002; Poorter et al., 2010; Choat et al., 2018). However, xylem transport is inherently vulnerable to hydraulic dysfunction during drought, as embolism formation within the xylem conduit network reduces hydraulic conductivity and can ultimately lead to canopy decline and tree mortality (Sperry & Pockman, 1993; Adams et al., 2017; Choat et al., 2018). Indeed, drought-induced hydraulic failure has been consistently identified as a proximate mechanism underlying tree mortality across taxa and biomes (Adams et al., 2017; Choat et al., 2012, 2018). Consequently, understanding how trees maintain xylem hydraulic integrity and which traits regulate proximity to hydraulic failure thresholds is critical to predicting drought resistance.

Trees use a spectrum of strategies to maintain functional xylem under drought (Klein et al., 2018). Embolism-avoidance strategies integrate mechanisms that limit water loss (e.g., early stomatal closure, leaf shedding) while enhancing internal storage, thereby preventing water potential from declining below critical thresholds that trigger xylem hydraulic failure. When such low water potential cannot be avoided, high resistance to xylem embolism (i.e., hydraulic safety) becomes essential for maintaining hydraulic function under drought stress. In contrast, some species rely on embolism-tolerance strategies that enable them to withstand higher levels of embolism. These strategies depend on rapid post-drought water uptake and the mobilization of non-structural carbohydrate reserves to restore hydraulic function through embolism repair or the production of new xylem (Brodersen & McElrone, 2013; Klein et al., 2018). Growing evidence suggests that the maintenance of xylem transport may be mechanistically coupled with multiple functional roles of bark (Rosell, 2019; Jupa et al., 2024).

Bark forms the interface between the stem and the atmosphere and comprises a structurally and morphologically variable complex of living and dead tissues that envelop the xylem (Angyalossy et al., 2016; Rosell, 2019). Anatomically, bark comprises all tissues external to the vascular cambium. The cambium produces secondary xylem inward and secondary phloem outward, representing the innermost part of the bark. Cortex is typically present above the phloem and contains mostly parenchyma cells produced by the primary ground meristem. The periderm is a secondary protective tissue produced by the phellogen, a secondary lateral meristem (Serra et al., 2022). Phellogen produces suberinized phellem (cork) outwards and parenchymatous phelloderm inwards. Phellem forms a functionally important protective barrier that limits water and gas fluxes and provides defense against pathogens (Lendzian, 2006). Due to its low permeability, phellem is locally modified by lenticels that enable gas exchange with the atmosphere (Rosner & Morris, 2022).

Bark performs several drought-relevant functional roles that can be mechanistically linked to xylem hydraulic limits. As the interface between the stem and the atmosphere, bark can substantially contribute to residual water loss when stomata close and leaf transpiration declines, thereby accelerating the decline in plant water potential. Bark water vapor conductance (G_bark_) varies widely among species and can be sufficiently high to induce stem water deficits and increase the risk of mortality from progressive embolism spread (Wolfe, 2020; Lintunen et al., 2021; Loram-Lourenço et al., 2022; Zhou et al., 2024). Under severe drought, water losses through bark may approach or even exceed leaf transpiration, highlighting the importance of bark in whole tree water balance (Lintunen et al., 2021). On the other side, water loss through permeable bark may be closely associated with its capacity to absorb water, a process that frequently occurs in both vapor (hygroscopic) and liquid phases (Mayr et al., 2014; Liu et al., 2019; Ilek et al., 2021; Losso et al., 2021; Gimeno et al., 2022).

Water that can be easily absorbed through the permeable bark can help to replenish internal water stores and may even be transported to the xylem, aiding embolism repair (Liu et al., 2019). Bark additionally serves as a critical reservoir for both water and carbohydrates. Internal water storage (capacitance) can buffer diurnal and drought-induced declines in water potential, thereby stabilizing hydraulic function and reducing the risk of failure (Meinzer et al., 2009; Scholz et al., 2011; Pfautsch et al., 2015; Bucci et al., 2023; Vega Grau et al., 2025). Simultaneously, carbohydrates stored in bark tissues or produced through bark photosynthesis can support the recovery of xylem transport capacity by sustaining cambial activity and facilitating the refilling of embolized conduits (Tomasella et al., 2021; Natale et al., 2023; Tian et al., 2025). Despite increasing interest in xylem-bark interactions, empirical evidence of their mechanistic integration remains limited, as the functions are typically investigated in isolation rather than within a unified functional framework.

Xylem and bark share deep developmental and evolutionary links, making the functional coordination between xylem hydraulics and bark traits biologically plausible. The true bark, as found in recent woody species, became evolutionarily prominent with the emergence of a vascular cambium and production of secondary xylem and phloem during the Early Devonian period, approximately 400 Mya (Morris & Jansen, 2017; Tomescu & Groover, 2019; Lalica & Tomescu, 2024). Fossil evidence further suggests that periderm-like protective tissues appeared early in the evolution of secondary growth, while the development of rhytidome became prominent later, coinciding with the rise of long-lived woody stems and the diversification of seed plants (Beck, 2010; Lalica & Tomescu, 2024). Over this shared evolutionary history, the coordination of xylem and bark functional properties was likely shaped by major environmental drivers governing plant performance, particularly water availability, temperature, and disturbance regimes. These selective pressures may have favored integrated trait syndromes linking hydraulic transport, protection, and storage functions, thereby promoting coordinated responses of xylem and bark to environmental stress.

Xylem efficiency and safety are shaped by climatic conditions that govern water availability, transpiration demand, and the frequency of hydraulic stress. Arid and climatically variable environments consistently select for embolism-resistant xylem, whereas humid and stable conditions favor higher transport efficiency (Gleason et al., 2016; Larter et al., 2017; Choat et al., 2018; Olson et al., 2020a; Grossiord et al., 2020; Liu et al., 2021; Lens et al., 2022). In contrast, ecological drivers of bark traits remain less resolved and appear to be governed by multiple, often interacting factors (Rosell et al., 2014; Rosell, 2019). Recent work suggests that bark water vapor conductance may respond to climatic gradients, increasing with mean annual precipitation and mean annual temperature across Neotropical species (Ávila Lovera & Winter, 2024). However, experimental stem heating in the same study produced the opposite response, underscoring the unresolved controls over bark conductance and related functional traits. Given that species with higher bark conductance experience greater water deficits and appear more prone to mortality from embolism (Wolfe, 2020; Loram-Lourenço et al., 2022), drought-prone environments may favor coordinated trait syndromes combining embolism-resistant xylem with bark that minimizes water loss, thereby strengthening embolism-avoidance strategies. Yet direct evidence for such integrated functional adaptation remains scarce.

Most studies examining links between xylem and bark rely on anatomical proxies rather than direct functional measurements (Poorter et al., 2014; Rosell et al., 2014; Jupa et al., 2024; McFarlane, 2024). As a result, we lack direct tests of (i) which bark functional traits align with xylem hydraulic strategies across species, (ii) whether these relationships are constrained by shared evolutionary history, and (iii) which climatic gradients drive coordinated adaptation in xylem hydraulics and bark function. This gap limits our mechanistic understanding of whole-tree drought resistance and hinders the integration of xylem–bark interactions into predictive frameworks of tree responses to drought.

Here, we address these gaps by testing functional coordination between xylem hydraulics and bark insulation and storage properties in juvenile stems of eight temperate woody Rosaceae species spanning contrasting ecological preferences and drought tolerance. We combine measurements of xylem transport efficiency and safety with key functional bark traits, including water vapor conductance, hygroscopic water exchange, water storage capacity, and anatomical investment in bark tissues.

We hypothesized that (i) xylem hydraulic and bark functional traits are coordinated across Rosaceae species, and ii) species from warmer and drier climates exhibit coordinated trait syndromes that jointly minimize dehydration and delay the onset of hydraulic failure (i.e., greater xylem embolism resistance associated with lower bark water vapor conductance and higher bark water storage).

## Materials and Methods

### Locality and plant material

For the experiment, we selected eight woody species from the Rosaceae family (three shrubs and five trees; Table **1**) growing in the area Kamenný vrch near the University Campus of the Masaryk University (Brno, Czech Republic; 49.1875633N, 16.5509383E, 370 m a.s.l., mean annual temperature 10.4°C, mean annual precipitation is 531 mm). The area is characterized by a scattered distribution of broadleaved and coniferous tree species growing on bedrock covered with Cambisol. From each species, we randomly selected 3-6 healthy mature individuals. The height of each tree individual was measured with TruPulse 200L Laser Rangefinder (LaserTech, USA), and the stem base diameter was measured 50 cm above ground with a caliper (Table **1**).

**Table 1.** List of species used in the experiments. Provided are the acronyms, growth form, the number of individuals used (n), mean height of the individual tree (height), mean stem base diameter (stem diameter), and mean age of the analyzed stem segments (age).

| Species | Acronym | Growth form | n | Height (m) | Stem diameter (cm) | Age (years) |
| --- | --- | --- | --- | --- | --- | --- |
| *Malus domestica* Borkh. | mado | tree | 5 | 4.3 | 18.6 | 3 |
| *Prunus avium* L. | prav | tree | 5 | 7.2 | 14.6 | 3 |
| *Prunus cerasifera* Ehrh. | prce | tree | 3 | 5.0 | 18.0 | 4 |
| *Prunus padus* L. | prpa | tree | 4 | 4.5 | 7.0 | 3 |
| *Prunus spinosa* L. | prsp | shrub | 6 | 3.3 | 3.7 | 3 |
| *Rosa canina* L. | roca | shrub | 5 | 2.6 | 3.6 | 3 |
| *Rubus fruticosus* L. | rufr | shrub | 4 | 3.1 | - | 2 |
| *Sorbus aucuparia* L. | soau | tree | 6 | 5.8 | 10.2 | 4 |

From each individual, we sampled one approximately 1.5-m-long, well-illuminated branch from the southern canopy. The branches were sampled on one day at the end of summer (August 24), during active growth in the early morning hours (6 – 8 am). All leaves and side branches were removed, and the branches were wrapped in black plastic bags with wet paper towels. The branches were immediately transported to the University of Innsbruck (Austria) for further processing. All the branches were stored cool until their processing.

### Branch processing

Each branch was cut into three segments, which were used for the particular analyses described below. The distal segment used for anatomical analyses was excised at distances of 25 cm and 30 cm from the branch tip and stored in FAA fixative after vacuum infiltration until further processing. The neighboring segment was excised at distances of 30 cm and 65 cm from the branch tip and used for hydraulic measurements. The proximal segment was excised at distances of 65 cm and 80 cm from the branch tip and used for assessing bark functional traits. The segments were excised at the same distances from the branch tip across species to minimize interspecific differences in segment age, which ranged from 2 to 4 years (Table **1**; Olson et al., 2020b).

### Anatomical analyses

For the anatomical analyses, permanent slides were prepared according to a slightly modified protocol described in Ruzin (1999). Briefly, the samples were dehydrated in a series of increasing butanol concentrations and embedded in paraffin (Paraplast, Leica, Germany). Subsequently, 15 µm-thick cross-sections were made with a sledge microtome, dewaxed in concentrated xylene and stained in 1% (w/v) Safranin dissolved in water. The sections were mounted on slides using Eukitt (Sigma-Aldrich) and allowed to harden for 48 h. The sections were then observed in bright field with an Olympus BX-50 microscope (Olympus, Japan). The section area was photographed using a Sony A35 camera (Sony, Japan) and merged into a single image in Adobe Photoshop (Adobe, USA). The xylem cross-sectional area as well as section radius and thicknesses of xylem (=distance between pith margin and mid of the cambial zone), secondary phloem (=distance between the mid of the cambial zone and the margin of the last ray cell, cortex (=distance between secondary phloem and phellogen), phellem (=distance between phellogen and last phellem cell) were measured in Fiji software (Schindelin et al., 2012). Total bark thickness included secondary phloem, cortex, and phellem thicknesses. In the branches studied, the phelloderm was poorly developed and indistinct from the cortex. For relevant comparisons, we standardized all thicknesses to the stem segment radius. Stem age was determined by counting the number of annual xylem rings.

### Xylem hydraulic measurements

The 35-cm-long segment used for hydraulic measurements was excised underwater. Approximately 10 cm wide stripe of the bark was removed from both segment ends with a sharp razor blade. Special attention was paid to not damage the xylem. The segment was shortened several times from both ends underwater with a sharp carving knife to release tension and avoid artefactual embolism (Torres-Ruiz et al., 2015), resulting in a final length of 27.4 cm. All native embolism was then removed by flushing the segment with distilled, filtered (0.22 µm) and degassed water enriched with 0.005 (v/v) Micropur solution (Katadyn Products, Wallisellen, Switzerland) under 100 kPa for 30 min. The xylem transport efficiency and safety were determined with the flow centrifuge (Cochard et al., 2005). The embolism-free segment was placed into a custom-built rotor in a Sorvall RC-5C centrifuge (Thermo Fisher Scientific, Waltham, USA). The samples were spun for 10 min at low RPMs corresponding to xylem water potential (Ψ) of -0.1 MPa. For the hydraulic measurements in the flow centrifuge, the same solution as for flushing of samples (see above) was used. Then, RPM was increased in 1 MPa steps to successively reduce Ψ until the percent loss of hydraulic conductivity (PLC, %) exceeded 95%. In each step, RPMs were held constant for 5 min before measuring the flow rate through the sample.

During the measurement, the PLC was calculated according to the following equation:

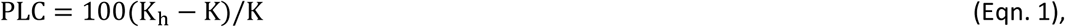

where K_h_ corresponds to the maximum hydraulic conductivity measured at Ψ=-0.5 MPa, and K is the actual hydraulic conductivity measured at a given Ψ.

Sigmoid vulnerability curves (plot of PLC vs. Ψ) were observed in all tested species, and the data were fitted with an exponential sigmoid function following Pammenter and Vander Willigen (1998):

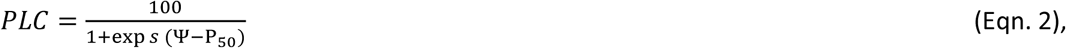

where *s* is a constant related to the curve slope, and P_50_ corresponds to the xylem water potential at 50% loss of hydraulic conductivity.

Xylem-specific hydraulic conductivity (K_xyl_; g m^−1^ MPa^−1^ s^−1^) was calculated as follows:

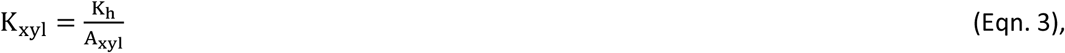

where A_xyl_ is xylem cross-sectional area.

### Bark functional properties

The proximal segment was used for measurement of bark water vapor conductance (G_bark_; mmol m^−2^ s^−1^), hygroscopic absorption time (H_time_; hours), actual hygroscopicity (H_act_; g cm^−3^), relative hygroscopicity (H_rel_), and maximum water storage capacity (C_bark_, g cm^−3^). Bark water vapor conductance was measured as described by Wolfe (2020). Initially, the 15-cm-long proximal branch segment was shortened to 10 cm on each side, and the segment was vacuum-infiltrated with water overnight. Both ends of the segment were sealed with Parafilm, and the precise dimensions of the segment were measured with a caliper (proximal and distal stem diameter, distance between Parafilm edges). The segments were dried with a paper towel to remove excess water, weighed on an analytical balance (Radwag AS 220.X2 PLUS, Radwag, Poland), and inserted into a thermostatically controlled cabinet (ET618, Lovibond, Germany), where the segments were kept at a stable temperature of 25°C and RH 55%. The segments were reweighed in regular 1 h intervals for 7 h in total. The temperature and RH were measured hourly using a digital thermal hygrometer (Voltcraft DL–141TH, Voltcraft, Germany) placed in the cabinet.

The bark water vapor conductance was calculated as:

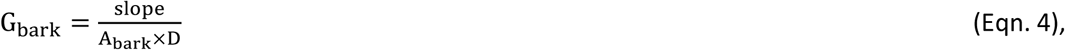

where slope (g s^−1^) represents the rate of water loss, A_bark_ (m^2^) represents the bark surface area of the segment, and D (dimensionless) is the vapor deficit. The slope was estimated using ordinary least squares, and only the linear part of the regression was retained (R^2^ > 0.98). The branch segment surface area was estimated as the lateral surface area of the truncated cone from the diameter and length measurements. Vapor deficit, i.e., the driving force of evaporation, was calculated as the mole fraction difference of water vapor in the stem (N_WStem_) and the surrounding air (N_WAir_) as:

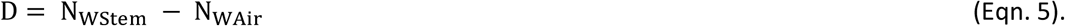

Both N_WStem_ and N_Wair_ were calculated according to the Buck equation (Buck 1981) as:

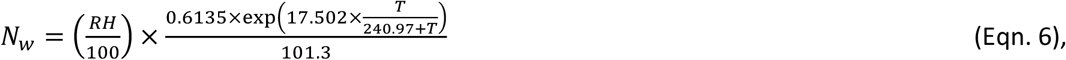

where RH (%) is relative humidity, T (°C) is the temperature, and 101.3 is the atmospheric pressure (kPa). The stem segments were assumed to be fully saturated by water (i.e., RH=100%) and in equilibrium with air temperature (Wolfe, 2020).

The bark hygroscopicity was determined according to the modified protocol of Ilek et al. (2016). The segment was infiltrated with water under vacuum overnight, and the bark was gently stripped from the segment. We paid special attention to remove an intact piece of bark from at least half of the branch circumference without damage. Only undamaged bark pieces were used for analyses. The bark volume (V_bark_; cm^3^) was then measured with the water displacement method. The bark segment was dried in a laboratory oven at 35°C to a constant weight (approx. 1 week). For larger bark segments, we stabilized the segment by inserting a silicone tube into the inner bark side to replace the xylem, thereby helping maintain the bark structure during dehydration and preventing warping or cracking. The hygroscopicity measurements consider water absorption only from the outer bark side. Therefore, a universal silicone was applied to the inner bark surface and allowed to dry overnight. The silicone weight was calculated as the difference in weight between the silicone-containing segment and the non-silicone segment.

The bark segments were inserted on a grid placed in a container partly filled with water, in which the relative air humidity reached 96%. The container with samples was placed in the thermostatically controlled cabinet and maintained at a stable air temperature of 25°C. The temperature and RH inside the box were regularly monitored with the Voltcraft digital thermal hygrometer. The segments were reweighed at regular intervals (typically once daily) until they reached a constant mass, a process that took 20-25 days.

The cumulative weight was plotted against time, and the data were fitted with a hyperbolic function:

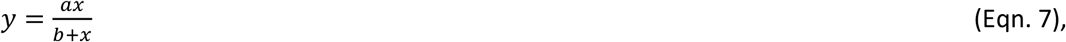

where a is the sample weight towards which the curve asymptotically grows and b is the time since the beginning of sample exposure when the increase of sample weight starts to level off. We defined the parameter b as hygroscopic absorption time (H_time_; hours). Greater H_time_ indicates slower water vapor uptake from the air, thus requiring more time to reach maximum hygroscopicity.

Subsequently, the segments were water-infiltrated under vacuum for 3 days, and the fully saturated samples were placed in a container at 100% relative humidity and left to drain for 24 h. After draining the loosely bound water, the samples were weighed to determine the saturated mass (m_s_), then dried to a constant weight at 70° C and reweighed to obtain the sample dry mass (m_d_). The maximum water storage capacity (C_bark_, g cm^−3^) was calculated as:

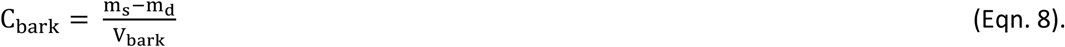

Actual hygroscopicity (H_act_, g cm^−3^), representing the maximum amount of water that can be absorbed by bark from air vapor, was calculated as:

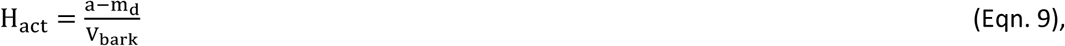

while, relative hygroscopicity (H*rel*) was calculated as the ratio of H_act_ and C_bark_.

### Phylogenetic analysis

Because species share evolutionary history, they cannot be considered statistically independent. To account for this non-independence and to quantify the degree of evolutionary conservatism in the traits studied, we reconstructed a phylogenetic tree for all examined species. Phylogenetic relationships were generated using the R package *U.PhyloMaker* (Jin & Qian, 2023), based on the *GBOTB.extended.WP.tre* megatree (Jin & Qian, 2022). Phylogenetic signal in individual traits was quantified using maximum-likelihood estimates of Pagel’s λ implemented in the *caper* R package (Orme et al., 2013). Pagel’s λ ranges from 0, indicating no phylogenetic signal, to 1, consistent with trait evolution under a Brownian motion model. The same maximum-likelihood framework was used to incorporate phylogenetic dependence when analyzing trait–trait and trait–environment relationships, as described below.

### Climatic data

The bioclimatic characterization of the species was derived from CHELSA v2.1 climatologies (1981–2010; Karger et al., 2017, 2021; Brun et al., 2022; downloaded on March 5, 2026) and species occurrences from the Global Biodiversity Information Facility (GBIF.org; downloaded on February 2, 2026). For each species, the occurrences were filtered by record type (HUMAN_OBSERVATION, OCCURRENCE, OBSERVATION, MACHINE_OBSERVATION, or PRESERVED_SPECIMEN), observation year (≥ 1900), coordinate uncertainty (≤ 5 km or missing), and by using CoordinateCleaner v. 3.0.1 (with the tests ‘capitals’, ‘centroids’, ‘gbif’, ‘institutions’, ‘seas’, and ‘zeros’; Zizka et al., 2019). Finally, the remaining occurrences were aggregated to the CHELSA grid, and climate values of grid cells with species presence were extracted and summarized by their median. This data has been deposited in ZENODO (reserved DOI: 10.5281/zenodo.20548945) and will be publicly available upon acceptance, together with a GBIF-derived dataset. Climate variables included mean annual near-surface air temperature (°C), annual precipitation (kg m⁻² yr⁻¹), mean annual vapor pressure deficit (Pa), and isothermality (a dimensionless index relating diurnal and annual temperature variability).

### Statistical analyses

To investigate multivariate coordination among the studied traits, we performed a principal component analysis (PCA) using the *FactoMineR* R package (Lê et al., 2008). Relationships among traits and between traits and environmental variables were first evaluated using linear regression based on species-mean values, with statistical significance assessed at α = 0.05. To account for phylogenetic non-independence among species, all relationships were subsequently re-analyzed using phylogenetic generalized least squares (PGLS) models, incorporating maximum-likelihood estimates of Pagel’s λ as implemented in the *caper* R package (Orme et al., 2013). We report the results of standard linear models and indicate changes after accounting for phylogenetic relatedness. In cases where the two approaches differed (indicated by Pagel’s λ > 0), we present estimates derived from the phylogenetic models to minimize potential bias associated with shared evolutionary history. All analyses were conducted in R version 4.5.1 (R Core Team; https://www.r-project.org).

## Results

### Interspecific variation in xylem and bark traits

The eight Rosaceae species differed in xylem hydraulics and in functional and anatomical traits of the bark (Table **S1**). Maximum hydraulic conductivity (Kh) ranged from 0.023 to 0.065 g m MPa^−1^ s^−1^, whereas embolism resistance was comparatively conserved, with the mean water potential causing 50% loss of conductivity (P_50_) spanning between −4.1 and −4.9 MPa. By contrast, bark functional traits varied widely across species, with bark water vapor conductance (G_bark_) ranging from 3.7 to 36.9 mmol m^−2^ s^−1^, hygroscopic absorption time (H_time_) from 7.7 to 43.2 hours, and maximum bark water storage capacity (C_bark_) from 0.58 to 0.80 g cm^−3^. The variability in bark functional traits was mirrored by substantial variability in bark anatomy (Table **S1**, Figure **S1**). In some species, phellem comprised numerous densely packed, thin-walled cells (typically in *Prunus* species) or fewer, thick-walled cells rich in phenolic compounds (typically in *Malus domestica, Rosa canina,* or *Sorbus aucuparia*; Figure **S1**). In *Rubus fruticosus*, unlike the other species, the phellem was developed in deeper cortical layers. In line, both absolute and standardized bark thickness (T_bark_: 582-1053 µm, TS_bark_: 0.15-0.28) and phellem thickness (T_phellem_: 18-121 µm, TS_phellem_: 0.005-0.033) substantially differed across species. Consequently, interspecific variation in most functional and anatomical bark traits exceeded that in xylem vulnerability to embolism (coefficients of variation: G_bark_ = 0.42, H_time_ = 0.60, C_bark_ = 0.10, TS_bark_ = 0.21, TS_phellem_ = 0.44 vs. P_50_ = 0.06).

### Trait relationships between xylem and bark

Xylem transport traits were closely linked to G_bark_ across species (Fig. **1a-b**). G_bark_ increased with both Kh and P_50_, indicating that species with higher xylem transport capacity and lower embolism resistance tend to have more permeable bark. Conversely, species with lower Kh and more negative P_50_ exhibited lower G_bark_. C_bark_ tended to be greater in species with lower Kh and P_50_, although these relationships were not statistically significant (Fig. **S2**). Other bark functional traits showed weak or no associations with xylem hydraulic traits (Fig. **S2**).

**Fig. 1.**
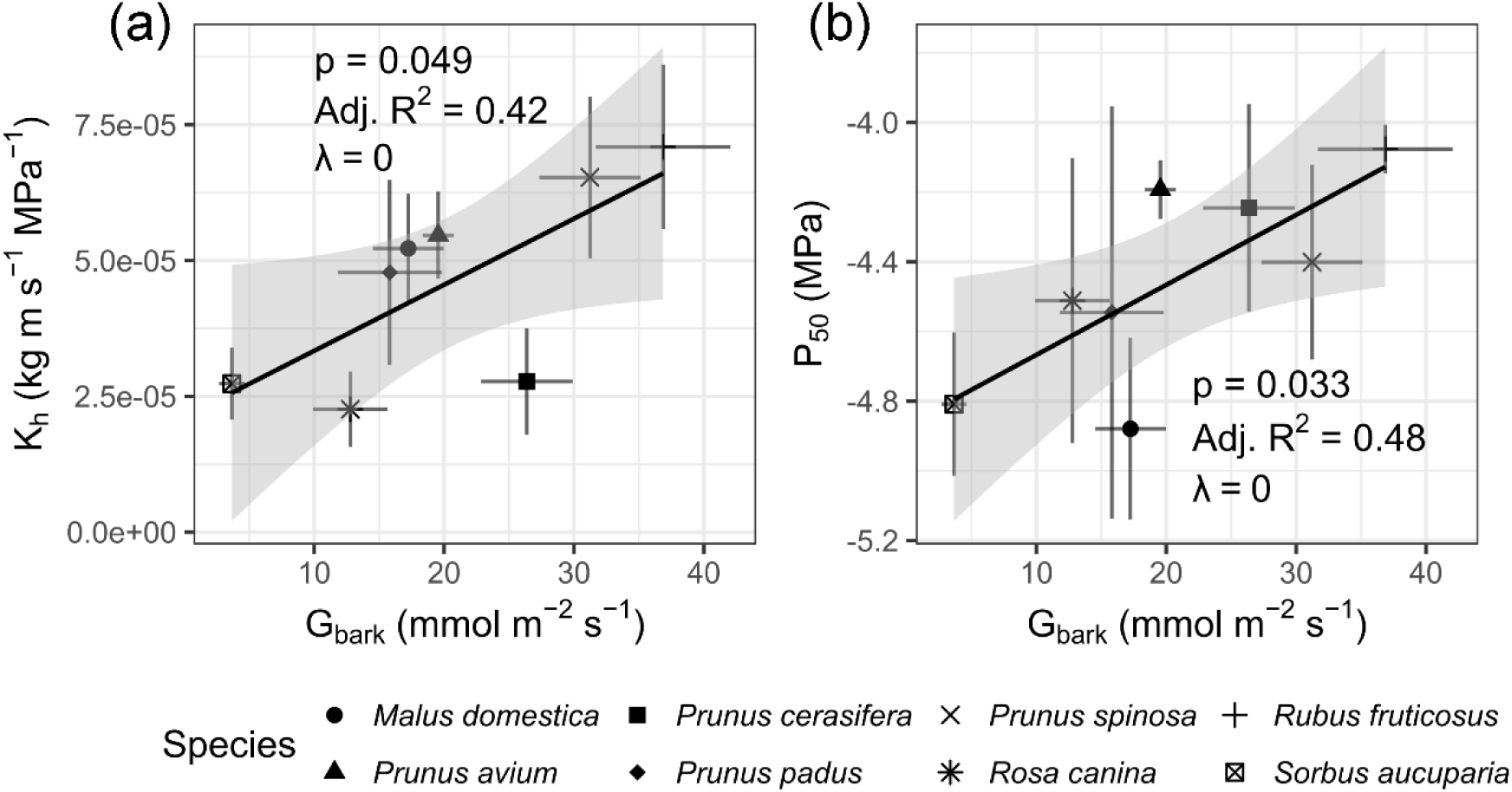
Relationships between bark water vapor conductance and xylem hydraulic traits across eight temperate Rosaceae species. Relationships of bark water vapor conductance (G_bark_, mmol m⁻² s⁻¹) with maximum hydraulic conductivity (Kh, kg m s⁻¹ MPa⁻¹; **a**) and xylem water potential at 50% loss of hydraulic conductivity (P_50_, MPa; **b**). Symbols represent species means, and error bars indicate ± SE. Solid lines show non-phylogenetic linear models with grey shaded areas indicating 95% confidence intervals. Statistical outputs (p-values, adjusted R²) and Pagel’s λ are shown within each panel. Across species, higher bark water vapor conductance was associated with greater xylem transport efficiency and less negative P_50_ values, indicating coordinated variation between bark permeability and xylem hydraulic strategy.

Besides, the correlation analyses further revealed mutual dependency of selected bark functional traits (Fig. **2**) and strong links of the bark functional traits to bark structure (Fig. **3**). Across species, G_bark_ declined with increasing H_time_ (Fig. **2**), highlighting a close functional coupling between water loss and hygroscopic water uptake. Species with less permeable bark required more time to replenish bark water reserves from atmospheric water vapor via hygroscopic processes. Both G_bark_ and bark hygroscopic properties were then primarily associated with TS_bark_ and TS_phellem_ (Fig. **3**). Specifically, G_bark_ was negatively related to TS_phellem_ (Fig. **3a**), indicating lower bark conductance in species with thicker phellem. A similar negative relationship was observed between G_bark_ and TS_bark_, although this association was marginally non-significant (Fig. **3b**). In contrast, the opposite relationships were observed for hygroscopic traits, when H_time_ was positively correlated with TS_phellem_ (Fig. **3c**) and TS_bark_ (Fig. **3d**). Similarly, TS_phellem_ correlated positively with H_act_ (Fig. **3e**) and H_rel_ (marginally insignificant correlation; Fig. **3f**). Functional relationships with both TS_bark_ and TS_phellem_ likely reflect the strong positive correlation between these two anatomical traits (Fig. **S2**), indicating coordinated anatomical variation within bark tissues.

**Fig. 2.**
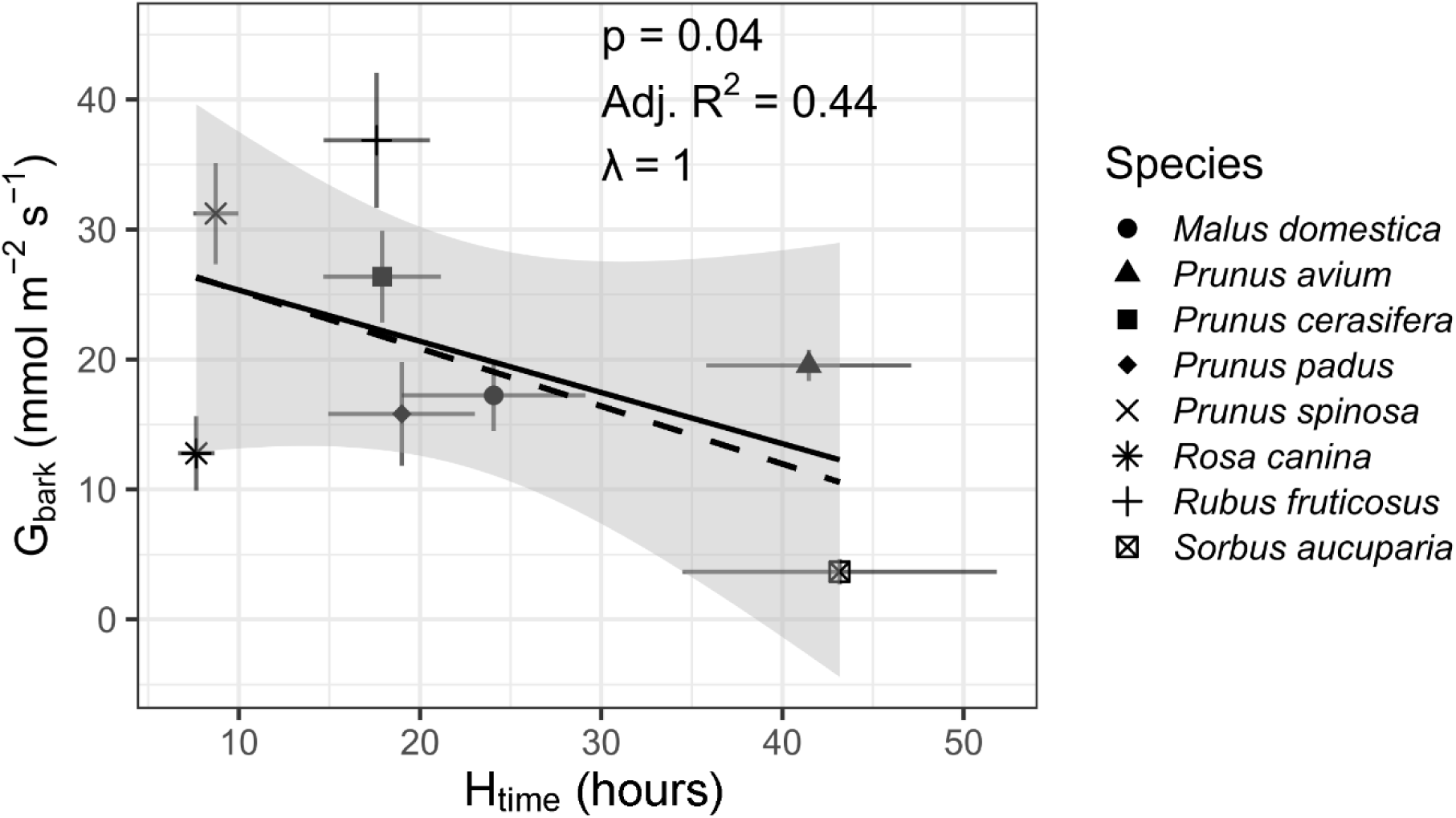
Relationship between bark water vapor conductance (G_bark_, mmol m⁻² s⁻¹) and hygroscopic absorption time (H_time_, hours) across eight temperate Rosaceae species. Symbols represent species means, and error bars indicate ± SE. Solid line indicates non-phylogenetic linear models with grey shaded areas indicating 95% confidence intervals; dashed line indicates fitted phylogenetic generalized least-squares (PGLS) relationships. Statistical outputs (p-values, adjusted R²) and Pagel’s λ are shown. Negative association between bark water vapor conductance and hygroscopic absorption time indicates that more slowly rehydrating bark is associated with reduced water vapor permeability.

**Fig. 3.**
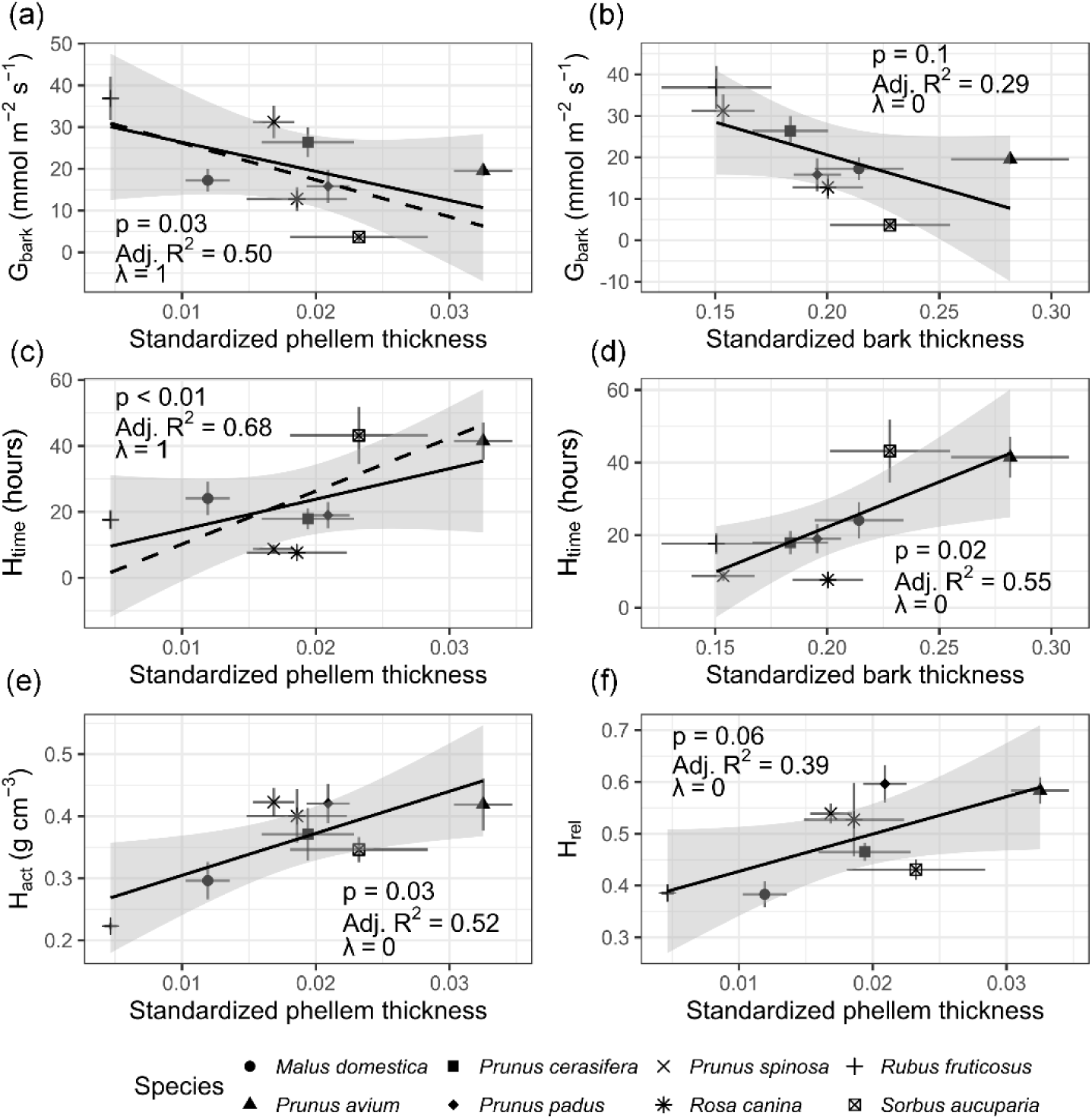
Relationships between bark functional properties and structural traits across eight temperate Rosaceae species. Relationships of bark water vapor conductance (G_bark_, mmol m⁻² s⁻¹) with standardized phellem thickness (**a**) and standardized total bark thickness (**b**), relationships of hygroscopic absorption time (H_time_, hours) with standardized phellem thickness (**c**) and standardized total bark thickness (**d**), relationship of actual hygroscopicity (H_act_, g cm⁻³) with standardized phellem thickness (**e**), and relationship of relative hygroscopicity (H_rel_) with standardized phellem thickness (**f**). Symbols represent species means, and error bars indicate ± SE. Solid lines show non-phylogenetic linear models with grey shaded areas indicating 95% confidence intervals; dashed lines indicate fitted phylogenetic generalized least-squares (PGLS) relationships, where applicable. Statistical outputs (p-values, adjusted R²) and Pagel’s λ are shown within each panel. Bark water-vapor conductance decreased with increasing phellem and total bark thickness, whereas hygroscopic properties showed the opposite tendency. This indicates that thicker bark tissues are characterized by lower permeability to water vapor, slower water-vapor uptake dynamics, but a greater capacity to store hygroscopically absorbed water.

### Phylogenetic patterns and associations with climate

The variability in functional and anatomical traits exhibited a strong phylogenetic signal across the studied species (Fig. **4**). Xylem vulnerability to embolism – P_50_ (Pagel’s λ = 1) and some of the bark functional traits – G_bark_ (λ = 0.7), H_act_ (λ = 0.9), and H_rel_ (λ = 0.7) showed strong phylogenetic conservatism. In contrast, Kh, H_time_, and standardized phellem and bark thickness exhibited little to no detectable phylogenetic signal (λ ≈ 0), with trait values varying substantially among closely related species, particularly within *Prunus* species.

**Fig. 4.**
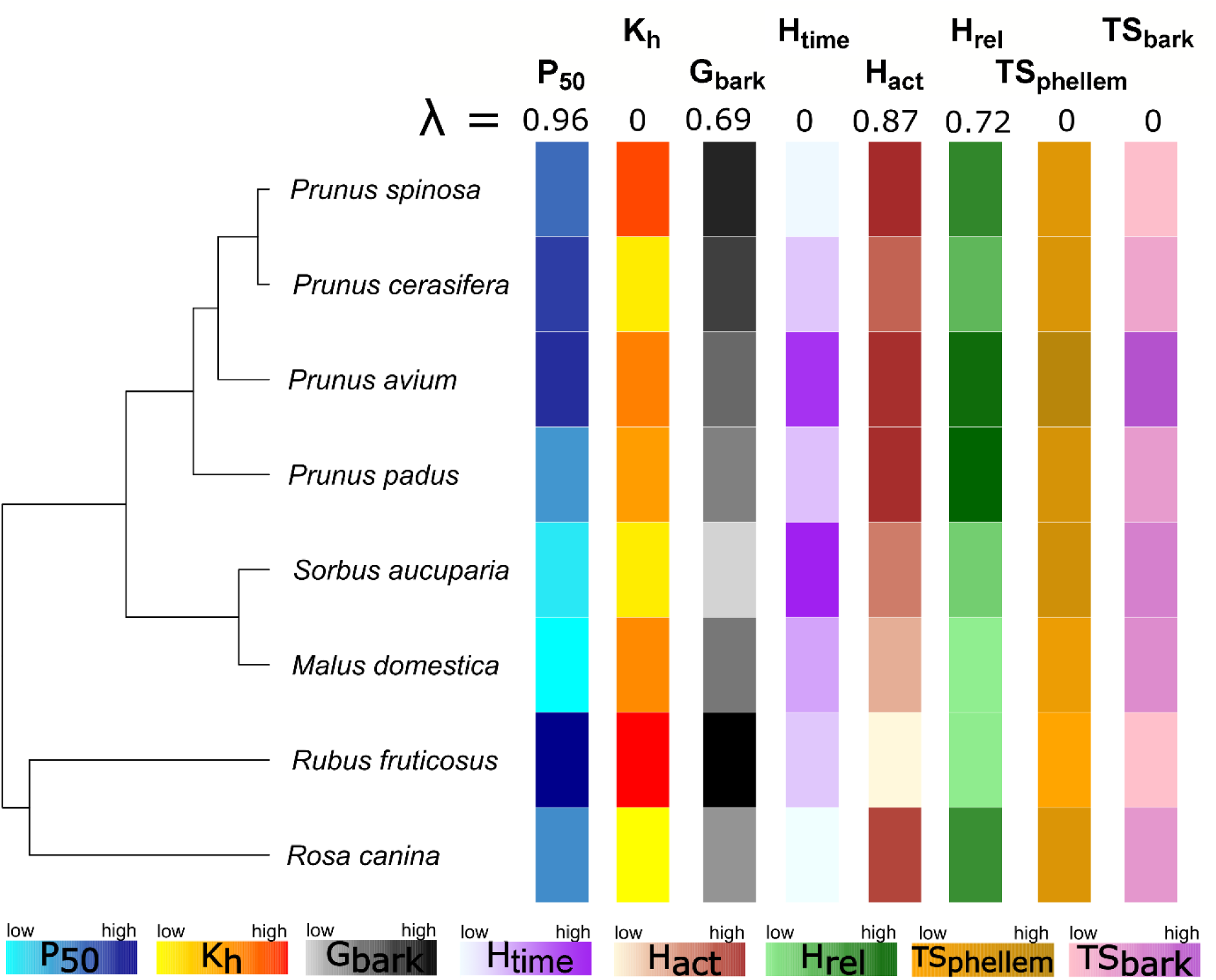
Phylogenetic distribution and trait variation across eight temperate Rosaceae species. Phylogenetic relationships among the studied species (left) and corresponding standardized trait values (right). Colored tiles represent relative trait magnitudes (from low to high) for xylem and bark functional and anatomical traits, including water potential at 50% loss of hydraulic conductivity (P_50_), maximum hydraulic conductivity (Kₕ), bark water vapor conductance (G_bark_), hygroscopic absorption time (H_time_), actual hygroscopicity (H_act_), relative hygroscopicity (H_rel_), standardized phellem thickness (TS_phellem_), and standardized total bark thickness (TS_bark_). Values are scaled within each trait to facilitate interspecific comparison. Pagel’s λ values for each trait are shown above the heatmap, indicating the strength of phylogenetic signal. Strong phylogenetic conservatism was observed for P_50_, G_bark_, and hygroscopic traits, whereas hydraulic conductivity and bark thickness traits exhibited weaker or negligible phylogenetic signal.

Among the functional traits studied, both P_50_ and G_bark_ showed strong dependence on multiple bioclimatic variables (Fig. **5**; Table **S2**). Both P_50_ and G_bark_ were primarily dependent on temperature-related variables, whereas precipitation-related variables showed generally weaker and mostly non-significant relationships (Table **S2**). Both P_50_ and G_bark_ concurrently increased with mean annual near-surface air temperature (Fig. **5a-b**) and mean annual vapor pressure deficit (Fig. **5c-d**). In addition, P_50_ and G_bark_ tended to increase with isothermality, suggesting that species with higher P_50_ and G_bark_ occur in environments with greater diurnal and lower seasonal variability in air temperature (Fig. **5e-f**). The remaining xylem- and bark-related traits showed weaker, nonsignificant relationships with the climatic variables (Table **S2**).

**Fig. 5.**
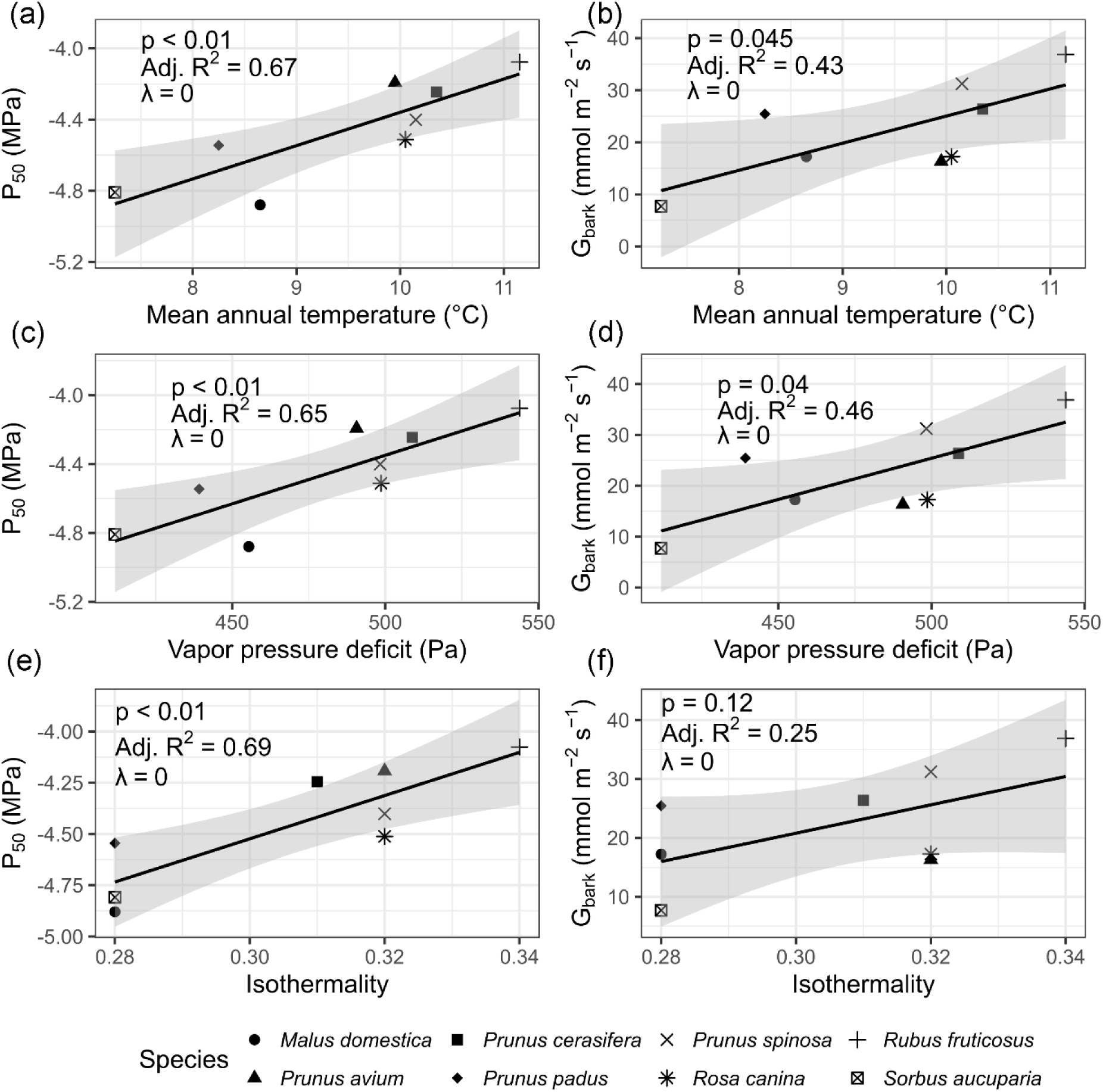
Climatic drivers of xylem vulnerability and bark water vapor conductance across eight temperate Rosaceae species. Relationships between xylem water potential at 50% loss of hydraulic conductivity (P₅₀; **a, c, e**) and bark water vapor conductance (G*bark*; **b, d, f**) with key climatic variables: mean annual near-surface air temperature (**a-b**), mean annual vapor pressure deficit (**c-d**), and isothermality (**e-f**). Solid lines indicate linear regressions with shaded 95% confidence intervals. Statistical outputs (p-values, adjusted R²) and Pagel’s λ are shown within each panel. Both P₅₀ and G*bark* increased with temperature, vapor pressure deficit, and isothermality, indicating coordinated shifts toward greater hydraulic vulnerability and higher bark permeability under warmer atmospheric conditions with greater diurnal variability in air temperature. Lack of phylogenetic signal (λ = 0) suggests that these relationships are primarily driven by environmental gradients rather than shared evolutionary history.

### Multivariate trait comparison

The principal component analysis (PCA) summarized the major trait covariation among species (Fig. **6**). The first principal component (PC1; 41% of the total variance) separated species preferentially occurring in warmer regions with higher vapor pressure deficit and greater isothermality (e.g., *Rubus fruticosus*, *Prunus spinosa*, *Rosa canina*, *Prunus cerasifera*) from species typically inhabiting rather cooler environments (e.g., *Prunus avium*, *Prunus padus*, *Sorbus aucuparia*). This climatic gradient broadly coincided with growth form, with shrubby species clustering towards warmer conditions and tree species associated with lower temperatures.

**Fig. 6.**
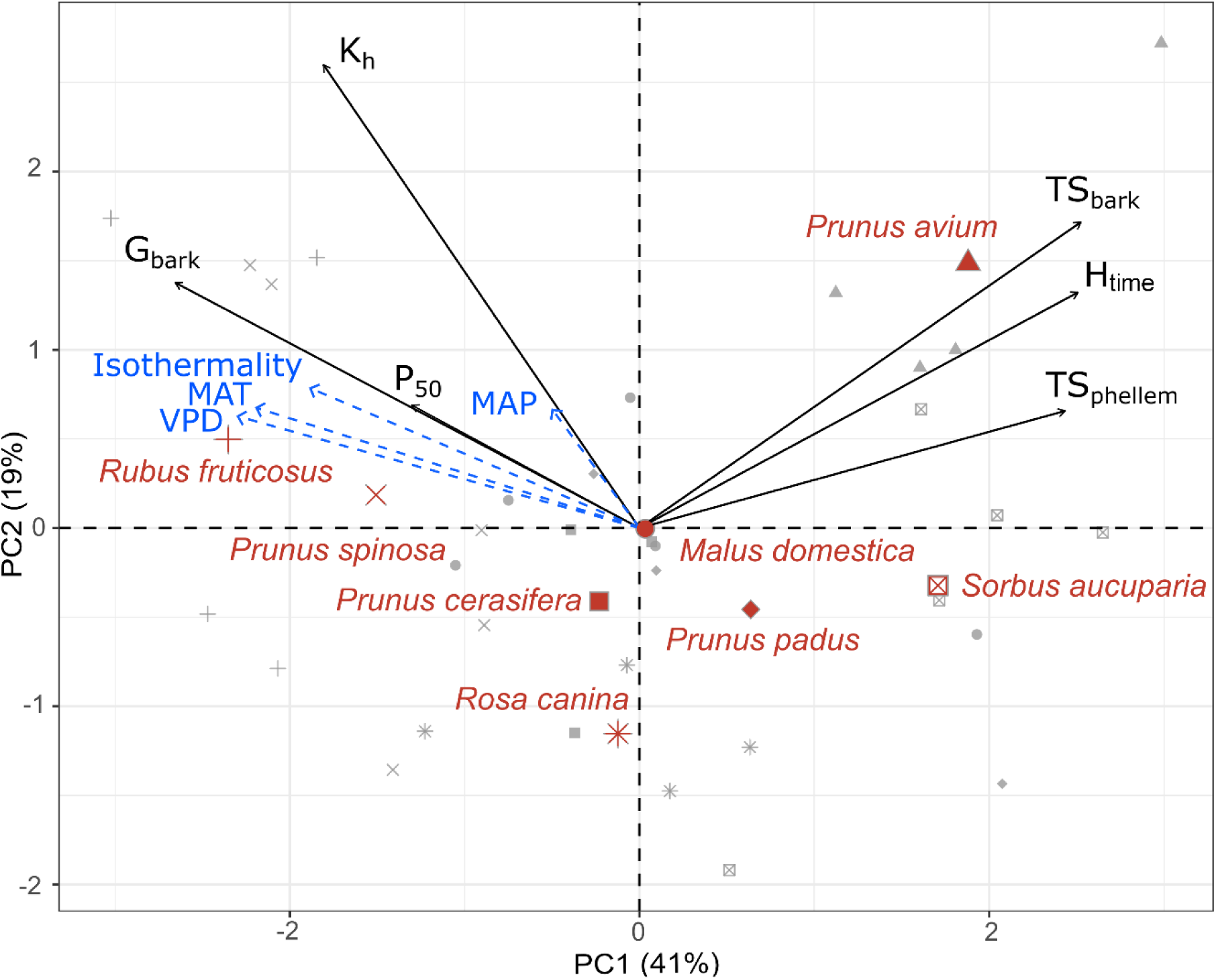
Principal component analysis of xylem and bark functional traits across eight Rosaceae species. Biplot of the first two principal components (PC1: 41%, PC2: 19%) summarizing trait covariation among species. Black arrows represent functional and anatomical trait loadings, including maximum hydraulic conductivity (Kh), xylem water potential at 50% loss of hydraulic conductivity (P_50_), bark water vapor conductance (G_bark_), hygroscopic absorption time (H_time_), and standardized bark and phellem thicknesses (TS_bark_, TS_phellem_). Blue arrows show climatic trait loadings, including isothermality, annual precipitation (MAP), mean annual near-surface air temperature (MAT), and mean annual vapor pressure deficit (VPD). Grey symbols indicate observations at the level of plant individuals. Species centroids are labeled red. PC1 captures a major gradient from species with high hydraulic efficiency, xylem vulnerability, and bark permeability (left) to those with greater bark thickness and slower hydration dynamics (right). This axis is strongly associated with temperature, indicating coordinated shifts in xylem and bark traits along climatic gradients.

Species associated with warmer climates were characterized by higher G_bark_, higher Kh, and less negative P_50_. In contrast, species from cooler regions exhibited higher H_time_ and developed thicker bark and phellem layers. Along with PC1, G_bark_ and P_50_ were closely related and aligned with isothermality, whereas H_time_ was primarily associated with TS_bark_ and TS_phellem_.

## Discussion

This study aimed to test whether bark functional traits and xylem hydraulic design are coordinated to optimize water relations and drought resistance in woody Rosaceae. In line with our expectations, xylem hydraulic efficiency and embolism resistance covaried with bark water vapor conductance and were closely associated with air temperature and atmospheric evaporative demands. These findings demonstrate that xylem and bark traits form an integrated functional system shaped by environmental constraints, underpinning species-specific drought response strategies. In the following sections, we interpret these relationships to elucidate how xylem–bark coordination contributes to whole-plant drought resistance.

### Functional xylem–bark interplay shapes drought resistance strategies

The association between xylem hydraulic traits and bark water vapor conductance indicates that interspecific variation in xylem water delivery is accompanied by coordinated variation in stem–atmosphere water exchange. More embolism-vulnerable Rosaceae species with higher xylem hydraulic efficiency tended to develop bark with high vapor permeability and lower water storage capacity (Fig. **1**, Fig. **S2**). High xylem transport efficiency facilitates water supply, which is important for tissue hydration and turgor maintenance (Cabon et al., 2020; Peters et al., 2021). Species with efficient xylem transport may thus rely more on rapid internal water supply and less on strong insulation against stem water loss, allowing for relatively high bark water vapor conductance. In this trait combination, fast xylem transport could partially compensate for greater bark-mediated water loss and help sustain stem water status under high stem–atmosphere conductance. Moreover, the permeable bark may simultaneously enhance O2 and CO2 exchange between the stem and the atmosphere, thereby supporting stem photosynthesis and respiration (Pfanz et al., 2002; Lendzian, 2006; Wittmann & Pfanz, 2014; Loram-Lourenço et al., 2022). Coordination between xylem transport efficiency and bark permeability may thus be central to maintaining high turgor and respiratory activity, which are key prerequisites for rapid stem growth in these species (Hölttä et al., 2010; Cabon et al., 2020).

Permeable bark may facilitate the rapid recovery of water reserves, which are essential for sustaining transpiration and buffering daily fluctuations in stem water potential (Meinzer et al., 2009; Steppe et al., 2015). Negative correlation between bark water vapor conductance and hygroscopic absorption time observed in our study clearly indicates that more conductive bark is capable of rapid atmospheric water uptake and prompt recovery of water reserves (Fig. **2**). The ability to rapidly replenish internal water storage via bark appears especially important for species with high xylem hydraulic conductivity, which can sustain high transpiration rates and therefore draw heavily on internal water reserves (Meinzer et al., 1995; Brodribb & Jordan, 2008).

The tight coordination between xylem hydraulic safety and bark permeability also appears to be an important mechanistic axis underlying the balance between embolism-avoidance and tolerance strategies. Water absorbed through permeable bark could be used directly for embolism repair (Liu et al., 2019; Beckett et al., 2023), while increased oxygen availability may sustain metabolically demanding processes that underlie xylem transport capacity, including cambial growth and the repair of embolized conduits (Brodersen et al., 2010; Secchi & Zwieniecki, 2011). Therefore, the potential of conductive bark for accelerated water absorption observed in more vulnerable Rosaceae species may compensate for generally greater demands of vulnerable xylem on frequent embolism repair (Brodersen et al., 2010; Choat et al., 2010). Conversely, reduced bark permeability and greater absolute water storage capacity in species with more embolism-resistant xylem likely limit water loss during drought and delay the decline of xylem water potential towards critical thresholds, inducing rapid embolism spread (Hacke et al., 2015; Skelton, 2020; Ruffault et al., 2022; Blackman et al., 2023).

### Phellem and bark thickness govern bark permeability and hydration dynamics

Results of this study identify phellem and total bark thickness as key traits shaping shifts between embolism avoidance and tolerance strategies in Rosaceae (Fig. **3**). Species with thicker bark consistently developed thicker phellem layers, reflecting coordinated activity of the vascular cambium and phellogen (Fig. **S2**). Such synchrony between secondary meristems has been documented in other woody taxa and likely reflects shared regulation by environmental cues and hormonal signals (Liphschitz, 1970; Angyalossy et al., 2016). Thicker bark enveloped by thicker phellem was characterized by lower bark water vapor conductance and prolonged hydration dynamics (Fig. **3a-d**). Functionally, thicker phellem acts as an effective diffusion barrier, slowing water exchange between the stem and the atmosphere, while thick bark mass appears to function as a water capacitor (Meinzer et al., 2004; Lendzian, 2006; Pfautsch et al., 2015). Due to its greater absolute water capacity, thicker bark can release larger volumes of water and thereby efficiently buffer a decline in stem water potential in more embolism-resistant species, but it requires a longer period to rehydrate (Meinzer et al., 2004; Pfautsch et al., 2015). In contrast, thin phellem combined with thin bark can result in poorer retention of acquired water, but shortened diffusion paths for water between the stem surface and xylem could increase the rehydration speed of both bark and xylem tissues in more vulnerable species.

Our findings on Rosaceae parallel recent evidence for coordinated evolution of bark structure and xylem hydraulic traits in Cupressaceae (Jupa et al., 2024), but the direction of some relationships differs. While embolism resistance was associated with thicker bark in both families, embolism-resistant Cupressaceae developed thinner phellem and thicker outer bark, unlike Rosaceae species. This contrast suggests that similar functional outcomes may be achieved through different bark architectures (e.g., rhytidome development, qualitative phellem traits, or lenticel density; Lendzian, 2006; Loram Lourenço et al., 2022; Rosner & Morris, 2022). In Rosaceae, phellem structure (many thin-walled cells or fewer thick-walled, phenolic-rich cells; Fig. **S1**) may increase diffusion resistance and partially compensate for the lack of early thick rhytidome that appears important for regulating bark water vapor conductance in conifers (Groh et al., 2002; Lendzian, 2006; Angyalossy et al., 2016; Jupa et al., 2024). Identifying determinants of bark conductance across gymnosperms and angiosperms is a key target for future research to understand species-specific drought resistance.

### Temperature and atmospheric evaporative demand shape xylem–bark coordination

Results of the phylogenetic analyses indicate that both bark water vapor conductance and xylem embolism vulnerability are phylogenetically conserved within Rosaceae, suggesting shared evolutionary histories of bark and xylem functional properties among the closely related lineages (Fig. **4**). The coordination between bark and xylem traits seems to be shaped by environmental conditions typical for species’ native habitats. Notably, among the mean climatic predictors considered, air temperature and atmospheric evaporative demand showed stronger associations with both bark and xylem functional traits than precipitation. To our knowledge, our results on Rosaceae provide the first evidence that atmospheric demand, particularly air temperature and VPD, may also drive the coordination between bark and xylem functioning.

Although the environmental drivers of bark water vapor conductance remain understudied compared to xylem hydraulics, emerging evidence supports the view that temperature and atmospheric evaporative demands can shape interspecific variation in bark permeability. Similar to our findings, a positive relationship between bark water vapor conductance and mean air temperature was revealed in a recent survey of 94 Neotropical woody species (Ávila-Lovera & Winter, 2024), reinforcing the idea that bark permeability is sensitive to climatic forcing.

However, in contrast to our expectations, Rosaceae species native to warmer regions combined higher bark water vapor conductance with lower xylem embolism resistance (Fig. **5**). This pattern in Rosaceae contrasts with the frequent finding across diverse woody lineages, where embolism resistance typically increases with aridity and climatic seasonality (Gleason et al., 2016; Larter et al., 2017; Choat et al., 2018; Olson et al., 2020a; Grossiord et al., 2020; Liu et al., 2021; Lens et al., 2022). One plausible explanation is that mean annual climate metrics may be poor proxies for the hydraulic stress actually experienced at species’ native sites. Drought intensity is determined not only by mean precipitation or temperature, but by their seasonal covariance, the timing of dry periods relative to phenology, and the frequency of extreme events over the season. Thus, the Rosaceae species associated with warmer, high-VPD climates do not necessarily experience stronger soil drought than species from cooler regions, particularly if precipitation seasonality or short-term extremes dominate stress episodes. Consistent with this interpretation, the Rosaceae species from warmer climates (*Prunus avium, Prunus spinosa, Rosa canina, Rubus fruticosus*) experienced, on average, higher annual precipitation than species from colder regions (*Malus domestica, Prunus padus, Sorbus aucuparia*; 10.3°C and 875 kg m^−2^ year^−1^ vs. 7.7°C and 806 kg m^−2^ year^−1^). These findings suggest that phylogenetically related temperate Rosaceae, occupying broadly comparable climates with generally modest chronic drought stress, may have evolved a VPD-filtered coordination between xylem and bark that prioritizes performance under recurring atmospheric demand.

In this sense, we propose that diurnal variability and short-term climatic extremes may impose stronger selective pressures on both bark and xylem traits than long-term climate means. This statement can be supported by the positive association of both P_50_ and G_bark_ with isothermality (index comparing day-night temperature variability with the annual temperature range; Fig. **5e-f**), suggesting that xylem vulnerability and bark conductance increase in environments with greater diurnal and lower seasonal variability in air temperature. Consistent with our results, climatic seasonality in precipitation and vapor pressure deficit has already been identified as an ecological filter shaping the co-optimization of xylem hydraulic efficiency and safety, with species from seasonally stable environments tending to build less hydraulically safe xylem (Liu et al., 2021). In contrast to xylem, bark, as the outermost layer, faces the atmosphere directly and may thus be directly affected by daily thermal fluctuations. This direct exposure to atmospheric forcing could also partially explain the higher interspecific plasticity observed in bark traits relative to xylem, whose structure may be more developmentally constrained and potentially more influenced by other environmental drivers, such as precipitation. Diurnal temperature oscillations and the accompanying diurnal fluctuations in atmospheric demand may thus represent an underappreciated axis of selection shaping coordination between xylem vulnerability and bark water vapor conductance.

In the Rosaceae studied here, the combination of higher bark water vapor conductance and lower xylem embolism resistance suggests a shift from conservative embolism avoidance toward embolism tolerance strategies in regions with more diurnally variable conditions and generally higher evaporative demands. In these regions, high diurnal VPD elevates internal water loss through increased transpiration and increases the risk of hydraulic failure, particularly in species with vulnerable xylem (Charrier et al., 2016; Schönbeck et al., 2022). Under this regime, higher bark permeability may be advantageous not by preventing embolism formation, but by promoting rapid growth recovery and embolism repair once drought stress is relieved, via facilitating water uptake through bark tissues (see above). Supporting this interpretation, our data on bark hygroscopic absorption time indicate that water uptake can be exceptionally fast in some species. Notably, the shrubby Rosaceae with the greatest bark water vapor conductance (*Rosa canina*, *Prunus spinosa*) recovered most of their bark water storage via hygroscopic rehydration within < 10 h. In regions with high diurnal changes in evaporative demand, this recovery potential may be constrained primarily to the nocturnal window of low VPD, when atmospheric conditions permit net rehydration. The functional coupling of hydraulically efficient yet embolism-vulnerable xylem with more permeable bark may thus help sustain growth and enhance drought resistance in climates characterized by diurnally variable conditions.

## Conclusions

Taken together, xylem and bark in Rosaceae species form an integrated functional system in which hydraulic safety, efficiency, and bark insulation are jointly tuned along diurnal and annual gradients of evaporative demand. Bark thus emerges not simply as a protective tissue, but as an essential component of whole-stem hydraulic function. This coordination supports a continuum of drought-response strategies, from conservative embolism avoidance to greater embolism tolerance, and indicates that drought resistance in woody angiosperms cannot be fully understood without considering bark traits alongside xylem hydraulics.

While this work substantially refines our understanding of drought resistance strategies in woody angiosperms, our test of the xylem-bark interaction was restricted to species from one family growing in a relatively narrow climatic gradient. Future comparative studies across wider phylogenetic and climatic ranges will be essential to determine whether xylem-bark coordination represents a general axis of functional evolution and to identify the climatic drivers of stem functional design. In these studies, incorporating short-term climatic extremes in addition to annual climate metrics will be important for resolving the climatic drivers of coordinated drought strategies.

## Acknowledgements

This work has been financially supported by the Czech Science Foundation (project no. 25-17559S).

## Author contributions

RJ designed the research, contributed to data analysis and interpretation, and wrote the original draft of the manuscript. TP, SM, and VG performed anatomical and physiological measurements and contributed to data analysis and interpretation. JD and JB conducted phylogenetic analyses and data evaluation. MN provided species occurrence and bioclimatic data. All authors reviewed and edited the manuscript.

## Data availability

All physiological and anatomical data supporting the results are available in the article and its Supporting Information (Figs **S1-S2**, Tables **S1-S2**). The bioclimatic data, together with a GBIF-derived dataset, have been deposited in ZENODO (reserved DOI: 10.5281/zenodo. 20548945) and will be publicly available upon acceptance.

