## Supplementary material for "The coordination between xylem and bark hydraulics in temperate Rosaceae species": Figure S1, S2, Table S1, S2

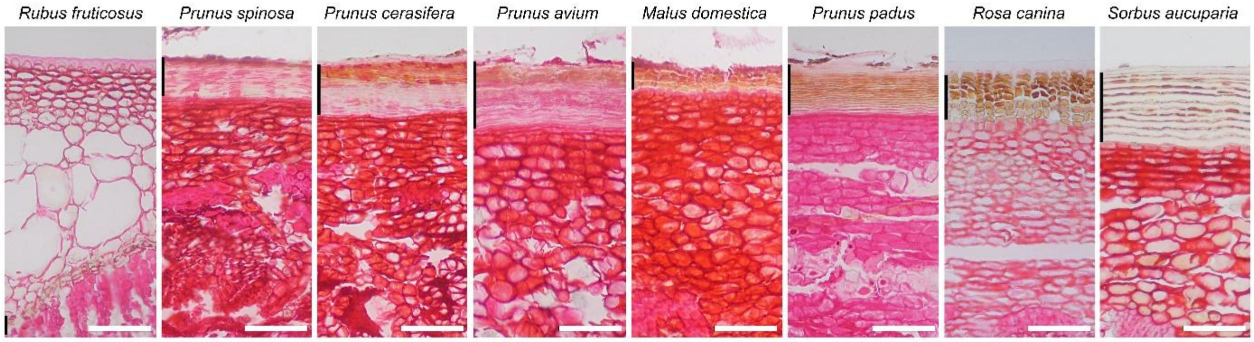


**Fig. S1. Transverse sections of the bark periphery of eight Rosaceae species.** Species are arranged from left to right according to decreasing bark water vapor conductance, with the highest values on the left and the lowest on the right. Sections were stained with safranin. The phellem layer is indicated by a black vertical bar along the left margin of each image. All images are shown at the same magnification. White scale bar = 100 µm.

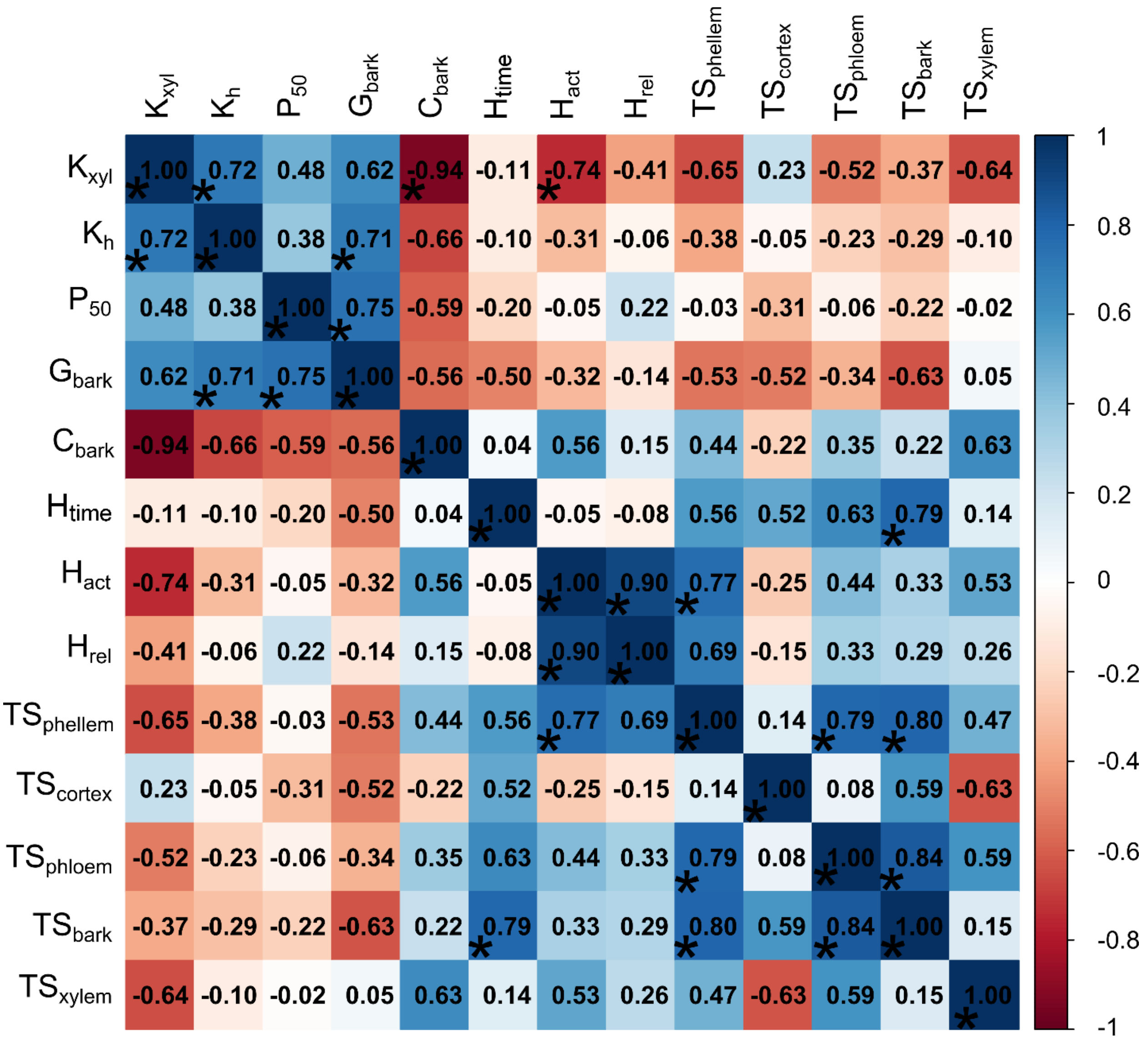


**Fig. S2. Correlation matrix of functional and anatomical traits related to xylem and bark of selected Rosaceae species.** Values represent Pearson correlation coefficients obtained for species means. Significant relationships ($P<0.05$) are highlighted by an asterisk. Legend: Xylem-specific hydraulic conductivity ($K_{xyl}$), maximum xylem hydraulic conductivity ($K_h$), water potential at 50% loss of hydraulic conductivity ($P_{50}$), bark water vapor conductance ($G_{bark}$), maximum bark water storage capacity ($C_{bark}$), hygroscopic absorption time ($H_{time}$), actual hygroscopicity ($H_{act}$), relative hygroscopicity ($H_{rel}$), standardized phellem thickness ($TS_{phellem}$), standardized cortex thickness ($TS_{cortex}$), standardized phloem thickness ($TS_{phloem}$), standardized bark thickness ($TS_{bark}$) and standardized xylem thickness ($TS_{xylem}$).

**Table S1. Comparison of functional and anatomical traits of xylem and bark across eight temperate Rosaceae species.** Data are presented in the form of mean ± SD (n=3-6). Functional and anatomical traits: Maximum xylem hydraulic conductivity ($K_h$), xylem-specific hydraulic conductivity ($K_{xyl}$), water potential at 50% loss of hydraulic conductivity ($P_{50}$), bark water vapor conductance ($G_{bark}$), maximum bark water storage capacity ($C_{bark}$), hygroscopic absorption time ($H_{time}$), actual hygroscopicity ($H_{act}$), relative hygroscopicity ($H_{rel}$), xylem thickness ($T_{xylem}$), bark thickness ($T_{bark}$), phloem thickness ($T_{phloem}$), cortex thickness ($T_{cortex}$), phellem thickness ($TS_{phellem}$), standardized xylem thickness ($TS_{xylem}$), standardized bark thickness ($TS_{bark}$), standardized phloem thickness ($TS_{phloem}$), standardized cortex thickness ($TS_{cortex}$) and standardized phellem thickness ($TS_{phellem}$). Species: *Malus domestica* (mado), *Prunus avium* (prav), *Prunus cerasifera* (prce), *Prunus padus* (prpa), *Prunus spinosa* (prsp), *Rosa canina* (roca), *Rubus fruticosus* (rufr), *Sorbus aucuparia* (soau).

| Parameter | Unit | mado | prav | prce | prpa | prsp | roca | rufr | soau |
|---|---|---|---|---|---|---|---|---|---|
| $K_h$ | g m s$^{-1}$ MPa$^{-1}$ | 0.052±0.022 | 0.055±0.018 | 0.028±0.017 | 0.048±0.034 | 0.065±0.037 | 0.023±0.016 | 0.071±0.030 | 0.027±0.016 |
| $K_{xyl}$ | kg m$^{-1}$ s$^{-1}$ MPa$^{-1}$ | 2.26±0.46 | 2.20±0.82 | 0.77±0.73 | 1.92±0.52 | 1.79±1.07 | 1.38±0.45 | 6.47±2.01 | 1.10±0.49 |
| $P_{50}$ | MPa | -4.88±0.58 | -4.19±0.19 | -4.25±0.52 | -4.54±1.18 | -4.40±0.69 | -4.51±0.91 | -4.08±0.14 | -4.81±0.50 |
| $G_{bark}$ | mmol m$^{-2}$ s$^{-1}$ | 17.24±6.10 | 19.55±2.70 | 26.36±6.10 | 15.82±8.00 | 31.22±9.55 | 12.78±6.40 | 36.86±10.39 | 3.67±2.40 |
| $C_{bark}$ | g cm$^{-3}$ | 0.77±0.07 | 0.71±0.09 | 0.79±0.11 | 0.71±0.13 | 0.79±0.11 | 0.78±0.14 | 0.58±0.02 | 0.80±0.09 |
| $H_{time}$ | Hours | 24.07±11.33 | 41.45±12.66 | 17.90±5.63 | 18.99±8.08 | 8.73±3.04 | 7.65±2.28 | 17.61±5.87 | 43.16±21.24 |
| $H_{act}$ | g cm$^{-3}$ | 0.30±0.07 | 0.42±0.11 | 0.37±0.07 | 0.42±0.06 | 0.42±0.06 | 0.40±0.10 | 0.22±0.01 | 0.35±0.05 |
| $H_{rel}$ | - | 0.38±0.06 | 0.58±0.06 | 0.46±0.03 | 0.60±0.07 | 0.54±0.05 | 0.53±0.16 | 0.39±0.01 | 0.43±0.05 |
| $T_{xylem}$ | µm | 1517±293 | 1891±372 | 2671±372 | 1402±812 | 2537±672 | 857±446 | 611±72 | 1554±599 |
| $T_{bark}$ | µm | 763±158 | 1053±211 | 771±203 | 677±128 | 609±91 | 684±156 | 582±192 | 826±248 |
| $T_{phloem}$ | µm | 349±151 | 510±170 | 441±201 | 292±72 | 260±61 | 245±144 | 173±53 | 319±130 |
| $T_{cortex}$ | µm | 387±38 | 434±119 | 242±29 | 304±43 | 284±39 | 377±13 | 402±147 | 421±134 |
| $T_{phellem}$ | µm | 42±12 | 121±13 | 79±15 | 74±23 | 64±13 | 61±21 | 18±3 | 80±26 |
| $TS_{xylem}$ | - | 0.421±0.046 | 0.501±0.064 | 0.646±0.091 | 0.371±0.151 | 0.603±0.160 | 0.234±0.062 | 0.158±0.021 | 0.417±0.127 |
| $TS_{bark}$ | - | 0.214±0.044 | 0.282±0.059 | 0.184±0.029 | 0.196±0.021 | 0.154±0.034 | 0.200±0.035 | 0.151±0.049 | 0.223±0.066 |
| $TS_{phloem}$ | - | 0.097±0.040 | 0.135±0.041 | 0.104±0.037 | 0.083±0.012 | 0.067±0.019 | 0.067±0.021 | 0.044±0.012 | 0.086±0.03 |
| $TS_{cortex}$ | - | 0.110±0.025 | 0.117±0.035 | 0.058±0.006 | 0.089±0.017 | 0.070±0.016 | 0.114±0.028 | 0.104±0.039 | 0.116±0.036 |
| $TS_{phellem}$ | - | 0.012±0.004 | 0.033±0.005 | 0.019±0.006 | 0.021±0.003 | 0.017±0.004 | 0.019±0.008 | 0.005±0.001 | 0.023±0.013 |

**Table S2. Relationships of functional (left) and anatomical traits (right) with selected bioclimatic variables.** Values represent Pearson correlation coefficients. Significant relationships ($P<0.05$) are highlighted in bold. Climatic variables: mean annual near-surface air temperature (temp), annual precipitation (prec), mean annual vapor pressure deficit (vpd), isothermality (isotherm). Functional and anatomical traits: Xylem-specific hydraulic conductivity ($K_{xyl}$), maximum xylem hydraulic conductivity ($K_h$), water potential at 50% loss of hydraulic conductivity ($P_{50}$), bark water vapor conductance ($G_{bark}$), maximum bark water storage capacity ($C_{bark}$), hygroscopic absorption time ($H_{time}$), actual hygroscopicity ($H_{act}$), relative hygroscopicity ($H_{rel}$), standardized phellem thickness ($TS_{phellem}$), standardized cortex thickness ($TS_{cortex}$), standardized phloem thickness ($TS_{phloem}$), standardized bark thickness ($TS_{bark}$) and standardized xylem thickness ($TS_{xylem}$).

| Functional trait | Climatic variable | r | Anatomical trait | Climatic variable | r |
|---|---|---|---|---|---|
| $K_{xyl}$ | Temp | 0.49 | $TS_{xylem}$ | Temp | -0.08 |
| $K_{xyl}$ | Prec | 0.52 | $TS_{xylem}$ | Prec | -0.48 |
| $K_{xyl}$ | Vpd | 0.55 | $TS_{xylem}$ | Vpd | -0.15 |
| $K_{xyl}$ | Isotherm | 0.54 | $TS_{xylem}$ | Isotherm | -0.20 |
| $K_h$ | Temp | 0.33 | $TS_{bark}$ | Temp | -0.41 |
| $K_h$ | Prec | 0.31 | $TS_{bark}$ | Prec | -0.05 |
| $K_h$ | Vpd | 0.34 | $TS_{bark}$ | Vpd | -0.45 |
| $K_h$ | Isotherm | 0.35 | $TS_{bark}$ | Isotherm | -0.32 |
| **$P_{50}$** | **Temp** | **0.84** | $TS_{phellem}$ | Temp | -0.34 |
| $P_{50}$ | Prec | 0.48 | $TS_{phellem}$ | Prec | -0.01 |
| **$P_{50}$** | **Vpd** | **0.84** | $TS_{phellem}$ | Vpd | -0.41 |
| **$P_{50}$** | **Isotherm** | **0.86** | $TS_{phellem}$ | Isotherm | -0.23 |
| **$G_{bark}$** | **Temp** | **0.72** | $TS_{cortex}$ | Temp | -0.32 |
| $G_{bark}$ | Prec | 0.15 | $TS_{cortex}$ | Prec | 0.38 |
| **$G_{bark}$** | **Vpd** | **0.73** | $TS_{cortex}$ | Vpd | -0.29 |
| **$G_{bark}$** | Isotherm | 0.60 | $TS_{cortex}$ | Isotherm | -0.11 |
| $H_{time}$ | Temp | -0.53 | $TS_{phloem}$ | Temp | -0.24 |
| $H_{time}$ | Prec | 0.12 | $TS_{phloem}$ | Prec | -0.39 |
| $H_{time}$ | Vpd | -0.52 | $TS_{phloem}$ | Vpd | -0.30 |
| $H_{time}$ | Isotherm | -0.37 | $TS_{phloem}$ | Isotherm | -0.31 |
| $H_{act}$ | Temp | -0.18 | | | |
| $H_{act}$ | Prec | -0.16 | | | |
| $H_{act}$ | Vpd | -0.28 | | | |
| $H_{act}$ | Isotherm | -0.16 | | | |
| $H_{rel}$ | Temp | -0.01 | | | |
| $H_{rel}$ | Prec | 0.05 | | | |
| $H_{rel}$ | Vpd | -0.10 | | | |
| $H_{rel}$ | Isotherm | 0.03 | | | |
| $C_{bark}$ | Temp | -0.47 | | | |
| $C_{bark}$ | Prec | -0.49 | | | |
| $C_{bark}$ | Vpd | -0.51 | | | |
| $C_{bark}$ | Isotherm | -0.50 | | | |
